# Adaptive evolution by spontaneous domain fusion and protein relocalisation

**DOI:** 10.1101/091744

**Authors:** Andrew D Farr, Paul B Rainey

## Abstract

Knowledge of adaptive processes encompasses understanding of the emergence of new genes. Computational analyses of genomes suggest that new genes can arise by domain swapping, however, empirical evidence has been lacking. Here we describe a set of nine independent deletion mutations that arose during the course of selection experiments with the bacterium *Pseudomonas fluorescens* in which the membrane-spanning domain of a fatty acid desaturase became translationally fused to a cytosolic di-guanylate cyclase (DGC) generating an adaptive phenotype. Detailed genetic analysis of one chimeric fusion protein showed that the DGC domain had become membrane-localised resulting in a new biological function. The relative ease by which this new gene arose along with its profound functional and regulatory effects provides a glimpse of mutational events and their consequences that are likely to play a significant role in the evolution of new genes.

## Introduction

The emergence of new genes by mutation - readily identified through comparative genomics - provides an obvious and important source of adaptive phenotypes (Chen et al. 2013; Long et al. 2013; Zhang and Long 2014). Mutational mechanisms involve divergence of duplicate genes (Ohno 1970; Lynch and Conery 2000; Bergthorsson et al. 2007; Nasvall et al. 2012), exon shuffling, the domestication of transposable elements, retrotransposition, gene fusion, and the de novo evolution of new reading frames (Long et al. 2003; Kaessmann 2010; Tautz and Domazet-Loso 2011; Ding et al. 2012; Long et al. 2013; Zhao et al. 2014).

Mutations that result in chimeric genes - by recombination of parental genes into a single open reading frame, or by retrotransposition of a gene into an alternate reading frame - are likely to generate new genes with relative ease (Rogers et al. 2010; Ranz and Parsch 2012). This is because fusions stand to produce novel combinations of functional domains and regulatory elements with few mutational steps. For example, promoter capture, whereby a fusion event couples an existing gene to a new promoter, can cause abrupt changes in temporal and spatial patterns of expression (Blount et al. 2012; Rogers and Hartl 2012; Annala et al. 2013). Additionally, novel combination of domains may result in a range of post-translational outcomes, ranging from relocalisation of domains, to novel inter-protein associations, regulation of enzymatic activity and possibly even the formation of novel protein functions (Patthy 2003; Bashton and Chothia 2007; Jin et al. 2009; Peisajovich et al. 2010; Rogers and Hartl 2012; Singh et al. 2012)

Comparative computational analyses provide evidence for the evolutionary importance of gene fusion events generating chimeric proteins. Compelling data comes a diverse range of organisms, including bacterial species (Pasek et al. 2006) *Oryza sativa* (Wang et al. 2006), *Drosophila* spp. (Long and Langley 1993; Wang et al. 2000; Wang et al. 2002; Jones et al. 2005; Rogers et al. 2009; Rogers and Hartl 2012), *Danio rerio* (Fu et al. 2010), *Caenorhabditis elegans* (Katju and Lynch 2006) and humans (Courseaux and Nahon 2001; Zhang et al. 2009). Despite the apparent importance of gene fusion events, the functional effects of presumed instances of domain swapping have received little attention. Notable exceptions are studies of “new” *Drosophila* genes: *sphinx, jingwei* and *Sdic*, which affect courting behaviour, alcohol-dehydrogenase substrate specificity and sperm-motility, respectively (Zhang et al. 2004; Dai et al. 2008; Zhang et al. 2010; Yeh et al. 2012). These three genes have complex mutational histories, but originate from gene fusion events: retrotransposition of distal genes (Long et al. 1999; Wang et al. 2002) or duplication and fusion of neighbouring genes (Ranz et al. 2003).

Beyond studies of new genes inferred from computational analyses, synthetic biology has shown that genes created by domain swapping can have important phenotypic effects. For example, in vitro recombination of genes involved in yeast mating was shown to generate greater diversity compared with control manipulations in which the same genes were duplicated in the absence of recombination (Peisajovich et al, 2010). While an elegant demonstration of the potential importance of gene fusion events in evolution, observation of such events in natural populations is desirable.

Evolution experiments with microbes provide as yet unrealised opportunities to understand evolutionary process, including mechanistic details. During the course of studies designed to elucidate the range of mutational pathways to a particular adaptive “wrinkly spreader” (WS) phenotype (Spiers et al. 2002; Goymer et al. 2006; Bantinaki et al. 2007; McDonald et al. 2009), we discovered a number of gene fusion events (Lind et al. 2015). Two classes bore the hallmarks of promoter capture whereby deletions caused a focal gene to come under control of a more highly expressed promoter eliciting the adaptive phenotype (Lind et al. 2015). A third class of mutation appeared not to conform to expectations of promoter capture. Intriguingly, this class was defined by eight independent deletions each of which caused a translational fusion between two adjacent genes. As we show here, these fusions define new genes that are chimeras between a membrane-localised fatty acid desaturase and a cytosolic di-guanylate cyclase (DGC).

## Results

### The WS model system and FWS types

When propagated in a spatially structured microcosm *P. fluorescens* SBW25 undergoes rapid diversification, producing a range of niche specialist types (Rainey and Travisano 1998). Among the best studied is the wrinkly spreader (WS) (Rainey and Travisano 1998; Spiers et al. 2002; Spiers et al. 2003; Goymer et al. 2006; Bantinaki et al. 2007; McDonald et al. 2009; Lind et al. 2015). WS types are caused by mutations in one of numerous genes that regulate or encode di-guanylate cyclases (DGCs). These genes catalyse the synthesis of 3', 5' -cyclic-di-guanosine monophosphate (c-di-GMP), a signalling molecule that allosterically regulates a complex of enzymes that produce an acetylated cellulosic polymer (ACP) (Ross et al. 1987; Tal et al. 1998; Amikam and Galperin 2006; De et al. 2008). Over-production of the polymer is the proximate cause of the WS phenotype (Spiers et al. 2002; Spiers et al. 2003).

Mutations that cause the WS phenotype in *P. fluorescens* SBW25 typically reside in one of three DGC encoding pathways: Wsp, Aws or Mws (McDonald et al. 2009). These pathways encode post-translational negative regulators that, when the targeted by loss-of-function mutations, result in constitutive DGC activity (Goymer et al. 2006). Elimination of these three pathways by deletion led to the discovery of a further 13 mutational routes to the WS phenotype, all involving pathways encoding DGC domains (Lind et al. 2015). Within this set of 12 new mutational routes were three loci where deletion events led to gene fusions. In two instances, the phenotypic effects were explained by increased transcription of DGCs resulting from promoter capture (Lind et al. 2015). However, for the third, which involved fusion between two open reading frames *(pflu0183* and *pflu0184)*, there was little to suggest that the phenotypic effects derived from promoter capture (Lind et al. 2015).

All mutations involving *pflu0183* described in Lind *et al.* (2015) involved deletions spanning parts of *pflu0183* and *pflu0184.* Each deletion resulted in a predicted single open reading frame, transcribed from the promoter of *pflu0184* (Figure 1A), thus maintaining the reading frame of *pflu0183.* Of the eight independent mutations at this locus, four consisted of an identical 467bp deletion (see Figure 1B and 1C). A ninth fusion, “L2-M5”, obtained from an independent experiment (see below) is also depicted in Figure 1. Reconstruction of the 467bp deletion in the ancestral SM genotype was sufficient to cause the WS colony morphology, niche specialisation and over-production of ACP (Figure 1 - figure supplement 1). All fusion mutations involving *pflu0183* produced the characteristic WS phenotype: hereafter these are referred to as “FWS” types and the focal FWS type studied here is termed “FWS2” (Figure 1 - figure supplement 2).

**Figure 1:**
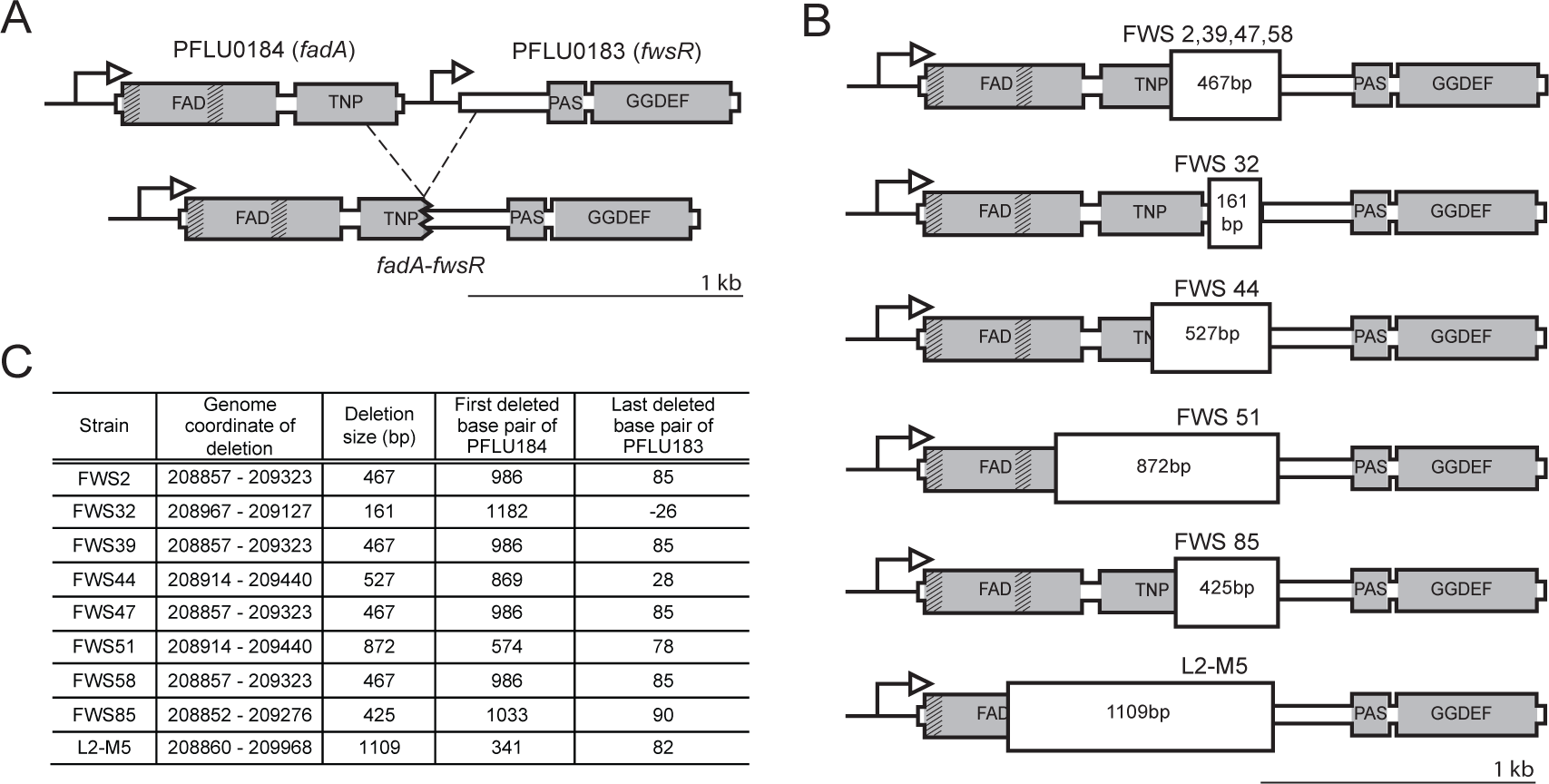
Arrangement of *pflu0184* and *pflu0183* in the ancestral SM genotype and following gene fusion. **A)** Illustrated is the fusion mutation that occurred on four independent occasions (underpinning FWS 2, 39, 47 and 58). Grey areas represent predicted domains and shading within grey areas depict predicted traosmemboane domains. *Pflu0184* (referred to as fndA) consists of a FAD region encoding a predicted fatty acid idesaturase at residues 11 to 246. ‘TNP’ is a predicted transposase element (DDE_Tnp_ISL3 family) at residues 261 to 389. The neighbouring *pflu0183* (referred to as *fwsR)* encodes a predicted PAS fold (of the PAS_3 family) at residues 68 to 157, and a predicted GGDEF domain at residues 174 to 331. The putative FadA protein contains two transmembrane domains (TMDs) between residues 10 to 32 and 135 to 157 as predicted by the ‘TMHMM server v2.0’ (Krogh et al. 2001). Dached lines represent the deletion that defines FWS 2, 3P, 47 and 58. **B)** Illustration of the nine independent *fadA-fwsR* fusions at *fadA* and f*WsR.* The mutations in FWS 2, 39, 47 and 58 share a common deletion junction. All deletions conserved the predicted ancestral reading frame for the GGEEF domain. **C)** Genomic details of *fadA-fwsR* fusion mutations. Positions are relative to the first base pair of the predicted open reading frame of *pflu0184* or *pflu0183*, or to the *Pseudomonas fluorescens* SBW25 genome (Silby et al. 2009).

### The *fadA-fwsR* fusion

*Pflu0183* encodes a predicted di-guanylate cyclase (DGC) henceforth referred to as ‘*fwsR’.* FwsR forms a predicted protein of 335 residues in length with a predicted PAS fold and a GGDEF domain at the C-terminus (Figure 1A). This GGDEF domain features a GGEEF motif, which is indicative of DGC activity (Chan et al. 2004; Galperin 2004; Malone et al. 2007; Wassmann et al. 2007). The neighbouring gene *pflu0184* encodes a predicted protein 394 residues in length, including a predicted N-terminal fatty acid desaturase (residues 11 and 246) and a predicted transposase element (residues 261 to 289). *Pflu0184* is hereafter referred to as *'fadA’.* FadA contains two predicted transmembrane domains (TMDs) between residues 10 to 32 and 135 to 157. The gene arising following deletion is termed *'fadA-fwsR'*.

Little is known about the functions of proteins encoded by either *fadA* or *fwsR.* FadA shares 83% identity with DesA (PA0286) from *Pseudomonas aeruginosa* (Winsor et al. 2011), which is a fatty acid desaturase (Zhu et al. 2006). Fatty acid desaturases modify phospholipids in the cell membrane in order to modify membrane fluidity in response to environmental change (Zhang and Rock 2008). In *P. aeruginosa*, transcription of *desA* is promoted during anaerobic conditions resulting in increased membrane fluidity as a consequence of double bonds introduced at the 9-position of fatty acid acyl chains (Zhu et al. 2006). Increased production of short-chained fatty acids has recently been shown to increase production of cyclic di-GMP by the Wsp pathway (Blanka et al. 2015). *fwsR* has orthologs across members of the genus *Pseudomonas*, however, none of these have been the subject of study.

It is not known whether the proximal relationship of *fadA* and *fwsR* orthologs across many *Pseudomonas* spp. reflects a functional or regulatory relationship between the two genes (or their protein products), however the operon prediction tool *DOOR* suggests they are separate transcriptional units (Mao et al. 2014). In species such as *P. putida* and *P. stutzeri* orthologs of *fadA* and *fwsR* are adjacent to each other (Figure 1 - figure supplement 3). However, in *P. aeruginosa* the locus of the GGDEF domain-encoding gene (PA0260) is separated from *desA* by three open reading frames (ORFs).

### DGC activity of FadA-FwsR is necessary to generate FWS2

How do deletions between *fadA* and *fwsR* (and generation of *fadA-fwsR)* produce FWS phenotypes? Like other WS-causing mutations (Goymer et al. 2006; Bantinaki et al. 2007; McDonald et al. 2009) the causal *fadA-fwsR* fusion is likely to alter the activity of the DGC domain of *fwsR* resulting in over-production of c-di-GMP. The constrained length of the spontaneous deletion mutations causing FWS types (all *fadA-fwsR* fusions arose by deletion of no more than 90 bp of *fwsR* - see Figure 1C) suggests that activity of the fwsR-encoded DGC is required for the FWS phenotype. To test this hypothesis the conserved GGEEF motif that defines the active site of the DGC was replaced by a mutant allele (GGAAF) expected to eliminate DGC function (Malone et al. 2007). The phenotype of FWS2 carrying the mutant *fadA-fwsR* allele was smooth, non-cellulose (ACP) producing and unable to colonise the air-liquid interface of static broth microcosms (Figure 2). This demonstrates that diguanylate cyclase activity of *fadA-fwsR* is necessary for the FWS2 phenotype.

**Figure 2:**
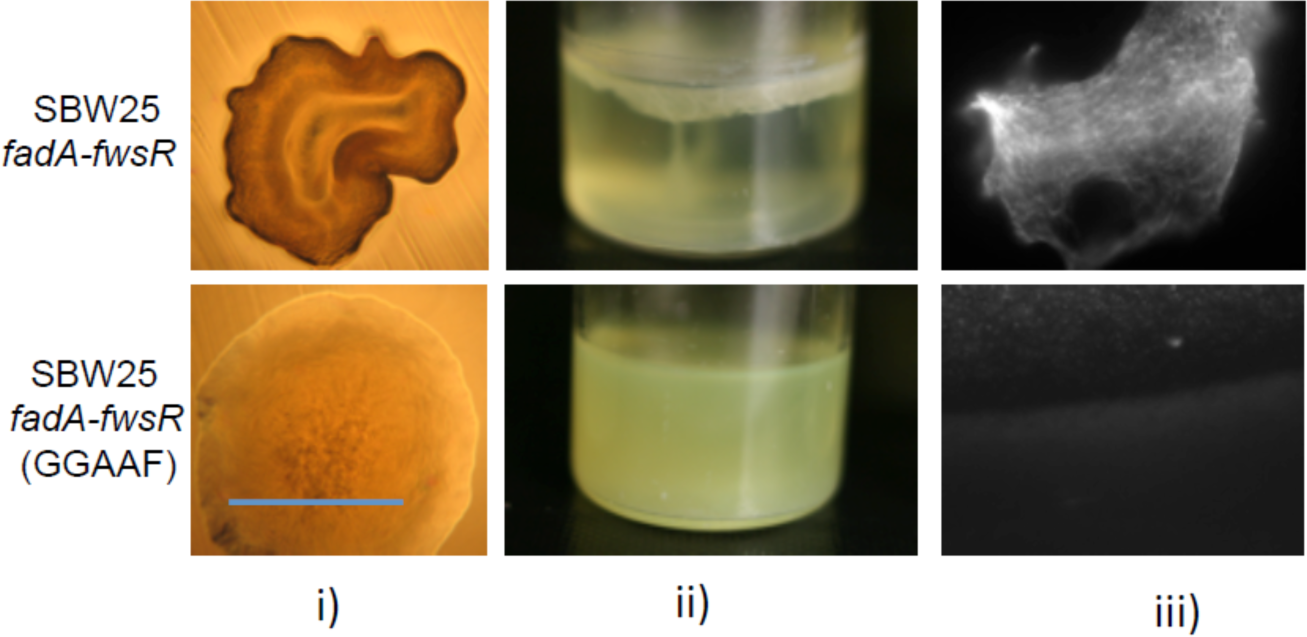
DGC activity encoded by *fadA-fwsR* is necessary to cause WS. In the context of *fadA-fwsR* reconstructed in SBW25, substitution of a GGAAF motif for the ancestral GGEEF motif within the predicted active site of the DGC domain results in SM phenotypes in i) morphology (visualised by light microscopy (10x objective) of ~32 h colonies grown on King’s B agar (KBA), blue bar is ~2mm); ii) colonised niche (depicted microcosms were inoculated with single colonies incubated statically at 28^o^C for 3 days)); and iii) ACP production, as indicted by calcofluor fluorescence (visualised by fluorescent microscopy, 60x objective).

### FWS2 does not require fatty acid desaturase activity

A *fadA-fwsR* fusion mutant isolated during the course of an independent experiment, termed ‘L2-M5’, contains a *fadA-fwsR* deletion that removed 347 bp of the predicted FadA domain (Figure 1B and 1C). This deletion results in a fusion of nucleotide 340 of *fadA* to nucleotide 83 of *fwsR*, while preserving the reading frame of *fwsR.* This ‘L2-M5’ mutation resulted in a characteristic WS phenotype upon substitution for *fadA* and *fwsR* in the ancestral SM genotype (Figure 3). That a *fadA-fwsR* fusion causes FWS, despite missing most of the enzymatic domain shows that fatty acid desaturase activity is not required for the FWS phenotype. It also suggests that structural componentry (such as non-enzymatic domain folds) of FadA is not required to cause FWS, unless that structure is proximal to the *N-* terminus.

**Figure 3:**
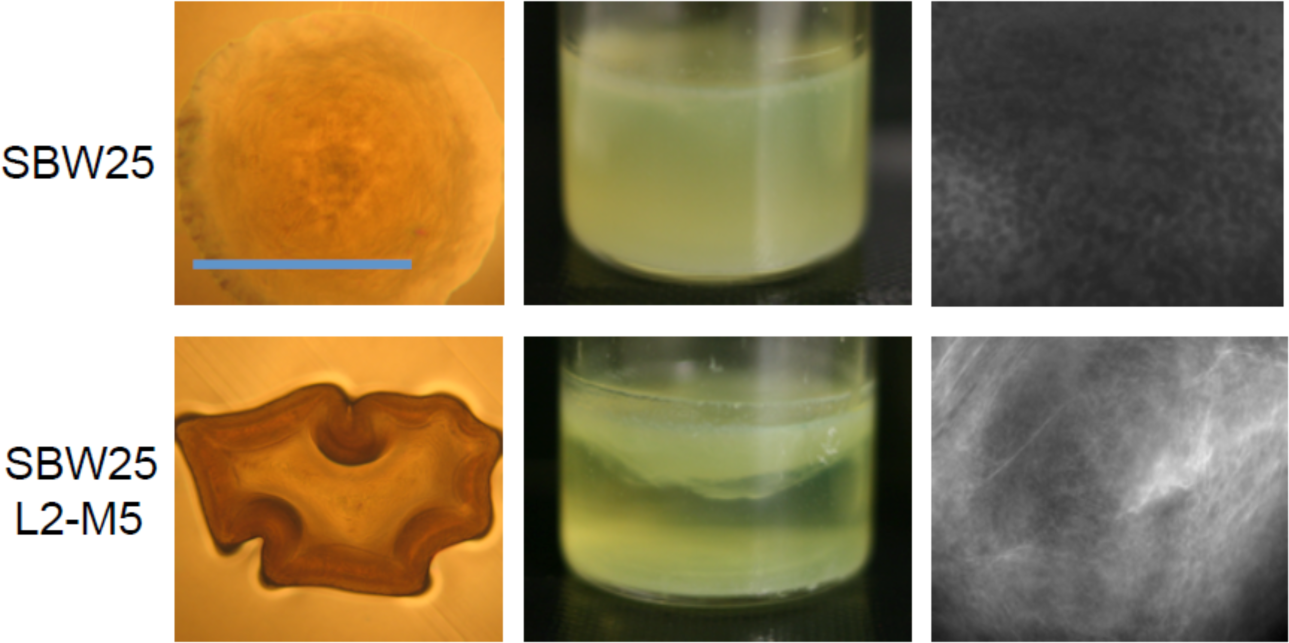
The fatty acid desaturase domain is not required for the phenotypic effects of *fadA-fwsR.* The phenotype of SBW25 expressing the L2-M5 *fadA-fwsR* fusion generates the characteristic WS phenotype, niche preference, and produces ACP as evidenced by presence of a fluorescent signal after staining a portion of the colony with calcofluor (see Figure 2 for details of each image).

### Altered transcription of *fwsR* is insufficient to cause FWS

One hypothesis to account for the effects of the *fadA-fwsR* fusion predicts effects mediated via transcription. Promoter mutations at several GGEEF domain-encoding genes have been associated with transcriptional increases and the evolution of WS types (Lind et al. 2015). In the case of the *fadA-fwsR* fusion, this possibility is given credence by the presence of a rho-independent transcriptional terminator downstream of the stop-codon of *fadA* (predicted by the WebGeSTer transcription terminator database (Mitra et al. 2011)). Removal of this terminator by the spontaneous deletion mutation could lead to increased transcription of *fwsR*, which may in turn elevate intracellular levels of the DGC, resulting in increased production of c-di-GMP and ultimately the FWS phenotype.

To test this hypothesis the promoter of *fadA* was fused to *fwsR* (resulting in *P*_*fadA*_*-fwsR*, Figure 4A) and integrated into the genome such that it replaced the *fadA fwsR* locus in the ancestral SM genotype. No effect of the *fadA* promoter fusion was observed: the genotype retained its smooth colony morphology, did not colonize the air-liquid interface, or produce ACP (Figure 4B). This demonstrates that changes in *fwsR* transcription caused by the *fadA-fwsR* fusion are not sufficient to cause WS.

**Figure 4:**
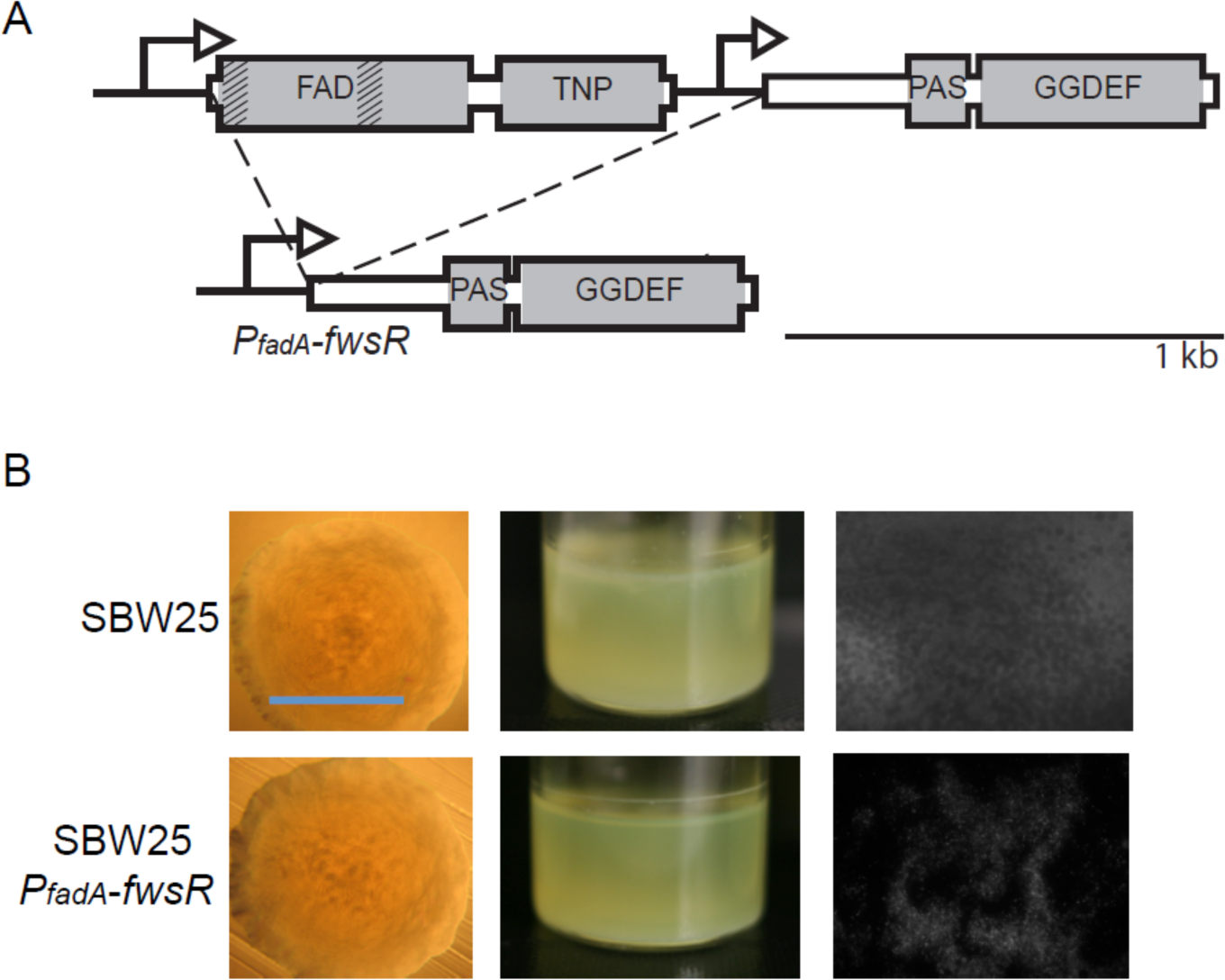
Altered transcription of *fwsR* is insufficient to cause FWS. **A)** Illustration of construct *P*_*fadA*_-*fwsR*. In this construct the promoter for fadA is fused to fwsR. **B)** The potential change in transcription caused by promoter capture does not cause the WS phenotype. The phenotype of the *P*_*fadA*_-*fwsR* construct reconstructed in the ancestral SM genotype is phenotypically SM in terms of morphology, niche specialization and absence of ACP production (see Figure 2 for details of each image).

### Translational fusion of FadA to FwsR is necessary for FWS2

The alternate hypothesis, namely, that the translation fusion is itself necessary to generate FWS2 was next tested. This was achieved by construction of a *fadA-fwsR* fusion in which translational coupling was eliminated: 182 bp between the stop codon of *fadA* and the ATG start codon of *fwsR* were deleted and replaced by stop codons (one in each reading frame). The *in situ* ribosome-binding site (RBS) of *fwsR* was left intact to allow translation initiation of an independent protein product for *fwsR.* The allele was introduced into the FWS2 background (where it replaced the *fadA-fwsR* fusion of FWS2) generating *fadA-3X-fwsR* (Figure 5A). The phenotype of this mutant resembled the ancestral SM type in all respects (Figure 5B). This indicates that translational fusion between *fadA* and *fwsR* is necessary to generate FWS2.

**Figure 5:**
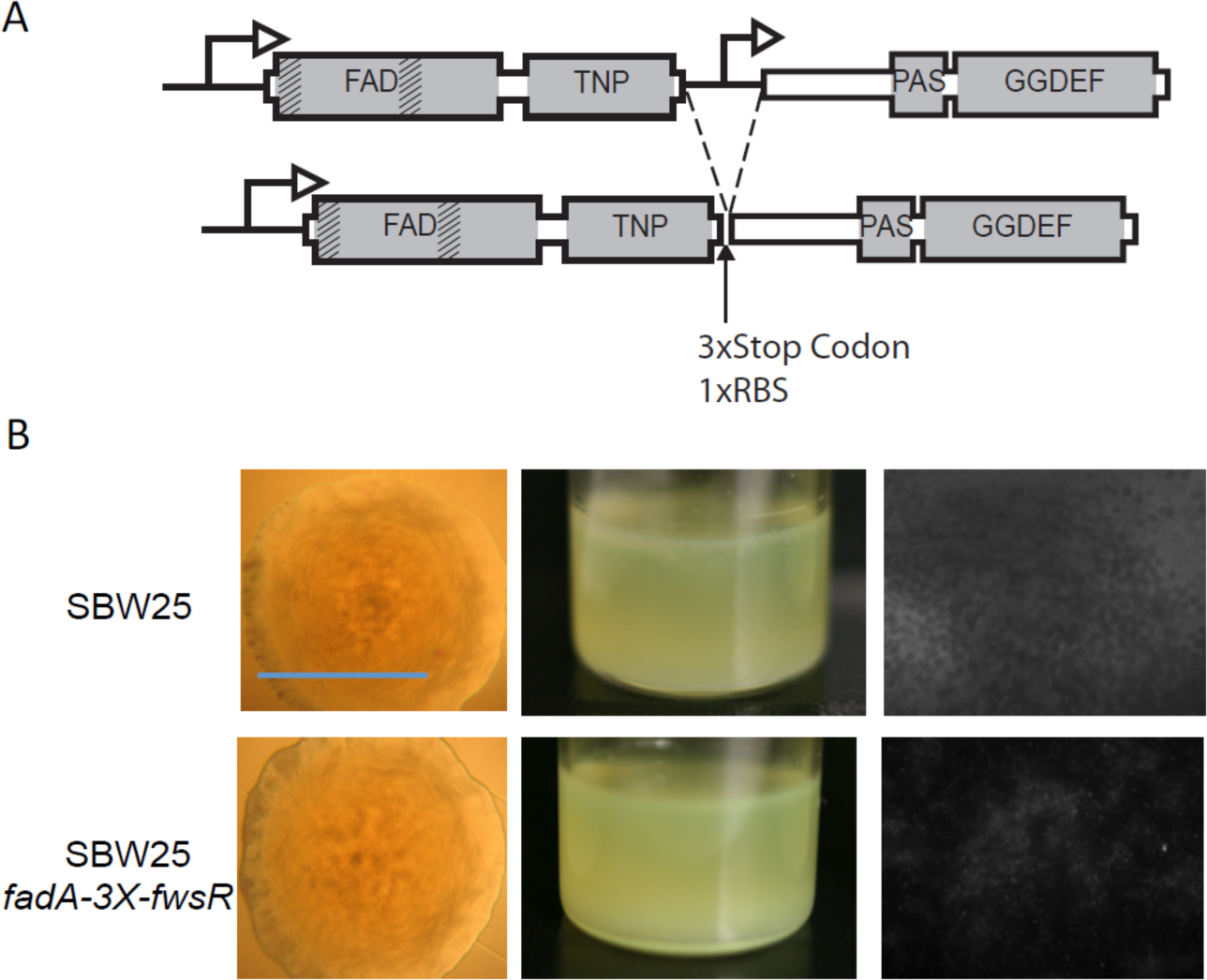
The FWS phenotype requires translational fusion of the *fadA* and *fwsR* genes. **A)** Illustration of the construct *fadA-3X-fwsR.* The *fadA* open reading frame was fused to that of *fwsR* and is separated by three stop codons. The ancestral ribosome-binding site (RBS) of *fwsR* was retained. **B)** Transcriptional fusion and translational uncoupling of *fadA* and *fwsR* do not cause FWS. The phenotype of *fadA-3X-fwsR* reconstructed in the ancestral SM genotype does not cause the FWS phenotype, indicating translational fusion of the genes is required to cause FWS. The *fadA-3X-fwsR* construct causes a characteristic SM phenotype in morphology, broth colonisation and lack of ACP production (see Figure 2 for details of each image).

### The FadA-FwsR fusion relocates the DGC domain

To test the hypothesis that *fadA-fwsR* results in localisation of FwsR to the membrane, the cellular locations of proteins encoded by *fadA, fwsR* and the *fadA-fwsR* fusion were visualised by creating a green fluorescent protein (GFP) translational fusion to the C-terminal region encoded by each gene. All fusions were cloned into the multiple cloning site (MCS) of mini-Tn*7-lac* and modified to contain a *Pseudomonas-specific* RBS (attempts to visualise foci using native gene promoters proved unsuccessful). Two controls were used during microscopy: a negative control of the ancestral SM genotype without a *gfp* insert in the mini-Tn7-*lac* cassette; and a positive control of the ancestral SM type containing *gfp* expressed in the integrated mini-Tn7-*lac* cassette. A cassette expressing *fadA-3X-fwsR-gfp* was also made to confirm that the *fadA-3X-fwsR* construct allowed independent translation of FwsR.

The fluorescent signal from each genotype was quantified to show that non-specific fluorescent signals from FwsR-GFP and FadA-3X-FwsR-GFP are significantly greater than the negative control (*Welch t*-test, *p*-value = 4.8 x 10^−10^ and 5.0 x 10^−8^), indicating that the diffuse expression of fluorescence results from the induced expression of GFP fusions (Figure 6).

**Figure 6:**
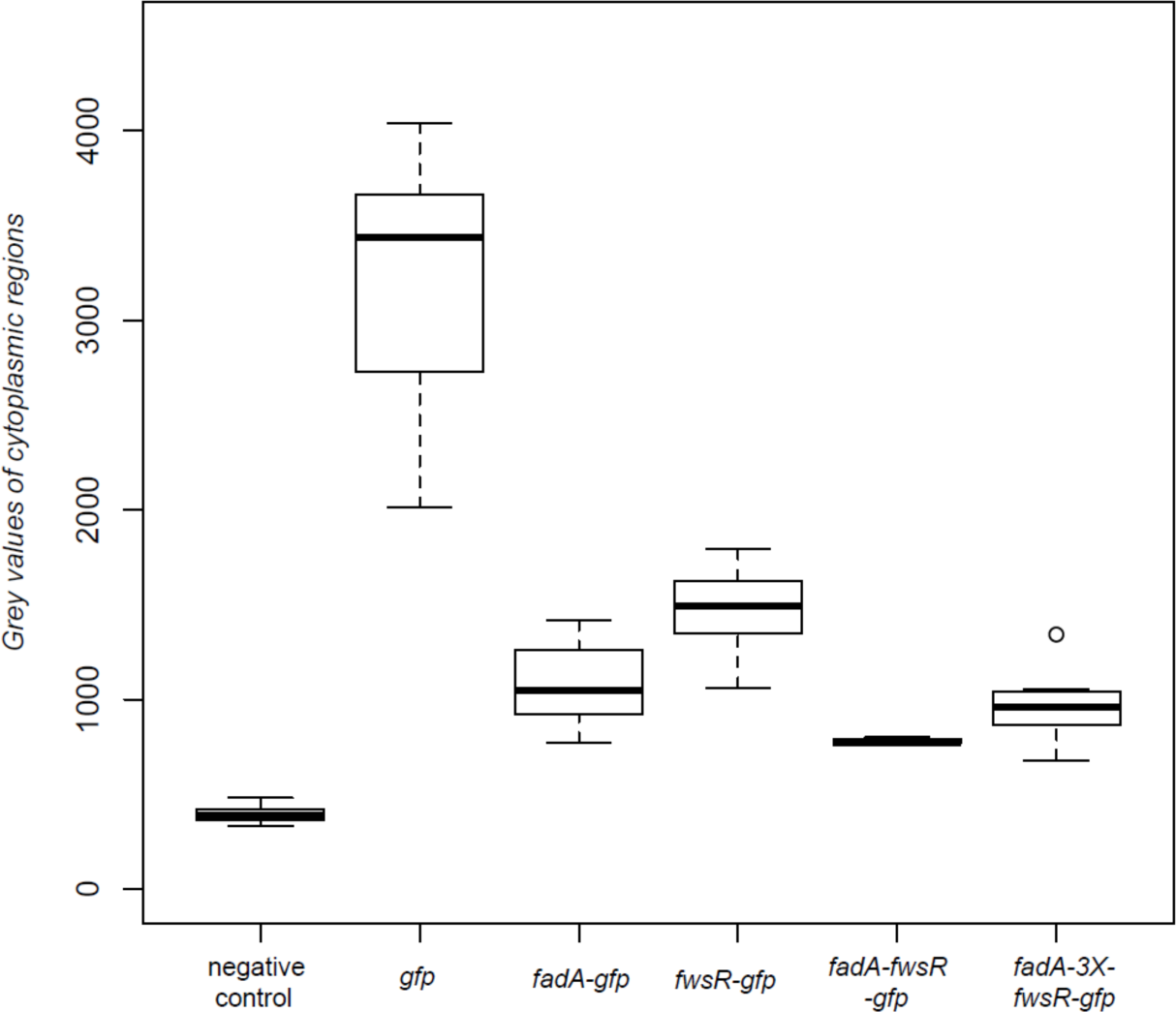
Cytoplasmic levels of fluorescence from protein fusions are above background levels. Depicted is a box plot of the florescent signals resulting from *gfp* translational fusions to fadA, fwsR and *fadA-fwsR* fusion variants. The mean fluorescent signal of 12 randomly picked cells (4 cells per independent culture) was obtained by making lateral transects of cells observed under fluorescent microscopy. The mean fluorescent signal of each cell was normalised to the background fluorescence of each image. Increases in the levels of fluorescence from the negative control demonstrate induced expression of genes fused with GFP. Data is presented as standard boxplots. Dark bars represent median values, boxes are the range from first to third quartile, whiskers represent minima and maxima, and circles represent values outside the 1.5 interquartile range.

Fluorescence microscopy demonstrates that the *fadA-fwsR* fusion alters the location of the FwsR DGC from the cytosol to the membrane (Figure 7). The location of the fluorescent signal of cells expressing *fwsR-gfp* is visually diffuse and is located within the cytoplasm (Figure 7C). Similarly, the signal from *fadA-3X-fwsR-gfp* is dispersed throughout the cytoplasm (Figure 7D). In contrast, cells expressing *fadA-gfp* have clear foci localised to the edge of the cell and, by inference, near the membrane (Figure 7E). The foci are distributed predominately at the lateral edge of cells in visual reference to the phase contrast images, similar to observations of the localisation of WspA in *P. aeruginosa* (O'Connor et al. 2012). The protein expressed by *fadA-fwsR-gfp* (Figure 7F) is localized in a manner identical to that observed in cells expressing *fadA-gfp.* The visual co-localisation of foci with the edge of the cells in genotypes expressing *fadA-gfp* and *fadA-fwsR-gfp* was confirmed by co-localisation analysis by using Van Steensel’s approach (van Steensel et al. 1996) (Figure 7 - figure supplement 1). Together, the location of fluorescent foci seen in cells with induced *fadA-gfp* and *fadA-fwsR-gfp* demonstrates the *fadA-fwsR* fusion event relocated FwsR from the native cytosol to the membrane location of FadA.

**Figure 7:**
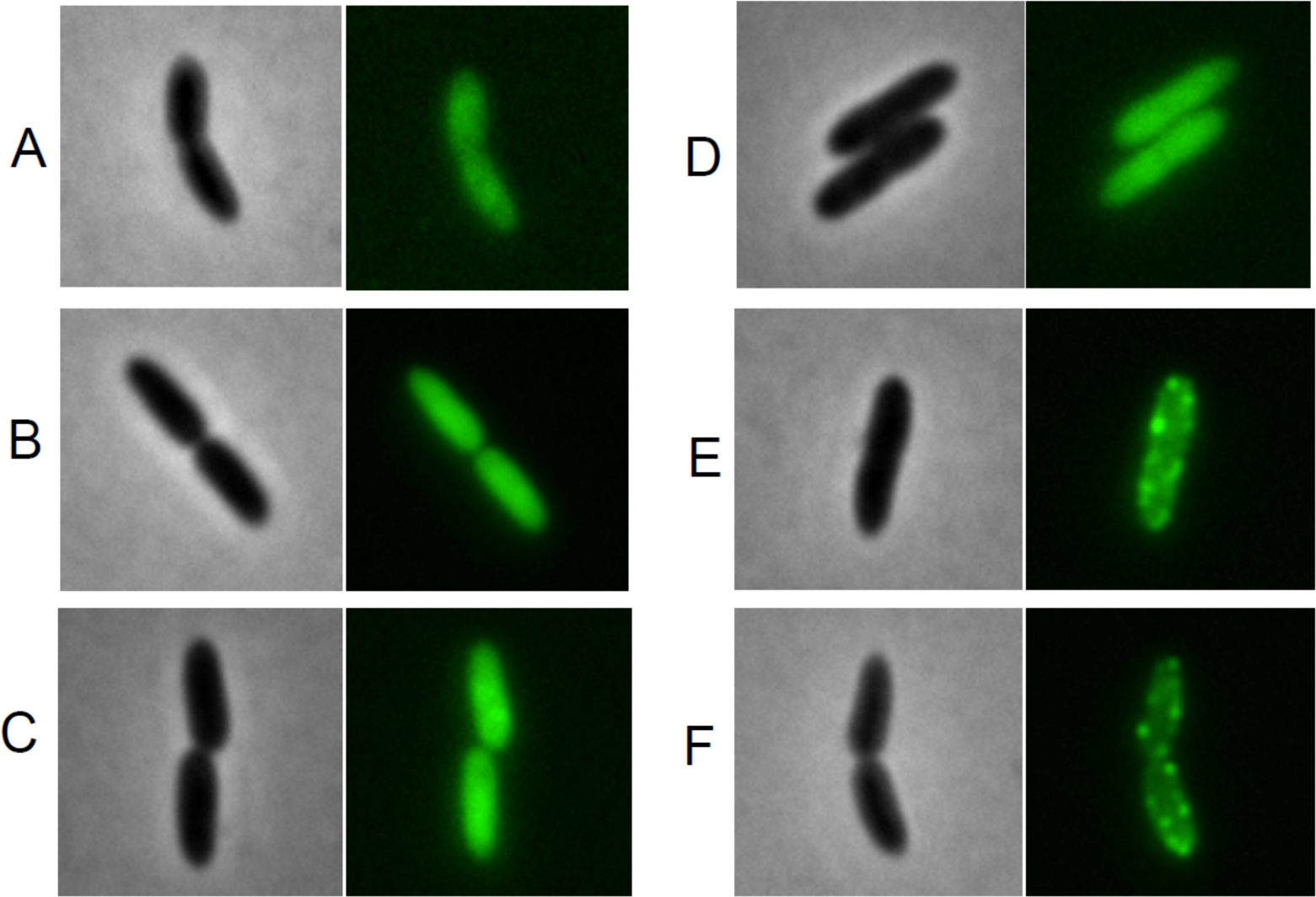
Fluorescence microscopy depicting the distribution of GFP tagged proteins encoded from the *fwsR* locus. Representative phase contrast (left) and GFP fluorescent images (right) of the subcellular distribution of fluorescence of A) *gfp(-)* control (showing background autofluorescence); B) gfp(+) control; C) *fwsR-gfp;* D) *fadA-3X-fwsR-gfp;* E) *fadA-gfp* and F) *fadA-fwsR-gfp.* Genotypes were grown in minimal media overnight and subcultured in minimal media with 1 mM IPTG for approximately 2 hours.

### Localisation of FwsR to the membrane is sufficient to generate FWS

If membrane localisation of FwsR is all that is required to activate synthesis of c-di-GMP, then fusion of FwsR to any membrane spanning domain-containing protein should suffice to generate the FWS phenotype. To test this hypothesis, the membrane-spanning domain of *mwsR* (PFLU5329) was fused to *fwsR.* Details of the construct are shown in Figure 8A. Replacement of the native *fadA fwsR* locus with the *mwsR-fwsR* fusion in the ancestral genotype lacking *mwsR* (to avoid unwanted allelic exchange at this gene) resulted in the FWS phenotype (Figure 8B).

**Figure 8:**
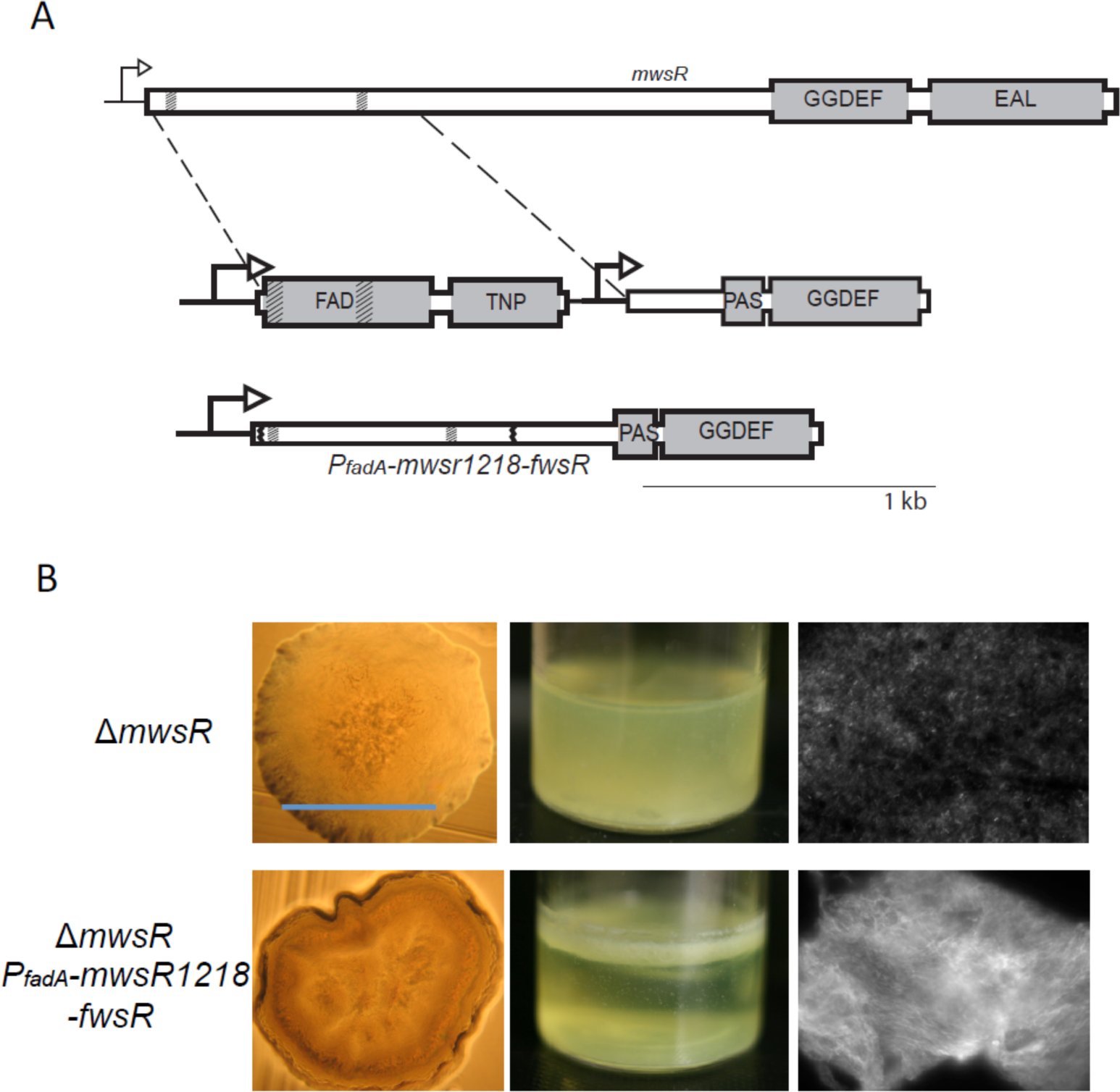
The translational fusion of the transmembrane regions of *mwsR* to *fwsR* causes the FWS phenotype. **A)** Illustration of the *P*_*fadA*_*-mwsR1218-fwsR* construct. The first 1218 bp of *mwsR* (which encodes for two TMD regions) was used to replace the *fadA* element of *fadA-fwsR.* Vertical grey bands within the ORFs represent TMDs. **B)** The phenotype of *PfadA-mwsR1218-fwsR* is FWS. The *P*_*fadA*_*-mwsR1218-fwsR* construct reconstructed in an SM ancestral background is characteristically WS in morphology, niche habitation and ACP production (see Figure 2 for details of each image).

## Discussion

The origin of new genes is a subject of fundamental importance and longstanding debate (Sturtevant 1925; Haldane 1932; Bridges 1936). Gene duplication and divergence, once seen as the primary source (Ohno 1970; Lynch and Conery 2000; Nasvall et al. 2012), is now recognized as just one of a number of routes by which new genes are born. Studies in comparative genomics indicate that new genes have arisen from retroduplication (in which mRNA is reverse transcribed into a complementary DNA copy and inserted into the chromosome (Brosius 1991; Long et al. 2003) from retrotransposon-mediated transduction (Moran et al. 1999; Cordaux and Batzer 2009), from deletion and recombination events that generate chimeric gene fusions (Marsh and Teichmann 2010; Ranz and Parsch 2012) from genomic parasites (Volff 2006; Feschotte and Pritham 2007), and even from previously non-coding DNA (Tautz & Domazet-Loso 2011, Neme & Tautz 2013).

Despite the apparent range of opportunities for the birth of new genes, there are few examples in which the evolution of a new gene has been captured in real time, the selective events leading to its formation understood, and mechanistic details underpinning function of the new gene revealed. Those thus far reported have involved recombination or deletion events leading to promoter capture. For example, in the Lenski long-term evolution experiment, Blount et al (2012) described a rare promoter capture event that underpinned evolution of citrate utilization in *E. coli.* Similarly, Lind et al (2015), using experimental *Pseudomonas* populations reported promoter capture events caused by deletions that increased transcription of genes encoding active DGCs necessary for over production of c-di-GMP and the evolution of WS types.

Here, provided with opportunities afforded by experimental evolution, we have observed, in real time, multiple independent deletion events, each of which caused a translational fusion between two genes: the membrane-spanning domain of *fadA* and the DGC domain of *fwsR.* Each fusion resulted in the birth of a new gene with the resulting fusion altering the cellular location of the DGC domain. This alteration conferred new biological properties: activation of the DGC, the synthesis of c-di-GMP, over-production of cellulose and generation of the adaptive wrinkly spreader phenotype.

In many respects, these events mirror those inferred from comparative studies responsible for new gene function, for example, the human gene *Kua-UEV* is thought to have originated from fusion of *Kua* and *UEV* resulting in localization of UEV to endomembranes by virtue of the N-terminal localization domain of *Kua* (Thomson et al. 2000). That similar, albeit simpler, events were detected in experimental bacterial populations is remarkable given that prokaryotic cell structure presents few opportunities for protein re-localization to occur; it thus adds weight to the suggestion that the birth of new genes via fusions that re-localize proteins is likely to be more common than recognized (Buljan et al. 2010). Indeed, inferences from comparative genomics support this prediction (Byun-McKay and Geeta 2007): approximately - 37% of duplicated gene pairs in *Saccharomyces cerevisiae* encode proteins that locate to separate cellular compartments (Marques et al. 2008). Of human multi-gene families known to encode proteins predicted to locate to the mitochondria, approximately 64% contain a gene predicted to relocate to an alternative subcellular location (Wang et al. 2009). Additionally, studies on the fate of duplicated genes in *C. elegans* suggest that approximately a third of new genes caused by duplication mutations are chimeric (Katju and Lynch 2003; Katju and Lynch 2006). Such chimeric mutations can introduce spatial encoding motifs to the duplicated protein, providing ample opportunity for the relocalisation of protein domains.

While the de novo fusion events reported here occurred between two adjacent loci, the fact that the DGC domain of *fwsR* could be fused (in vitro) to the membrane spanning domain of a gene more than one million nucleotides away *(mwsR)* and with similar effect reveals the considerable potential to generate new gene functions via fusion of distal genetic elements. This realization warrants consideration in context of standard models for the origin of new genes via duplication (amplification) and divergence (Ohno 1970, Bergthorsson et al 2007, Lynch and Conery 2000, Nävsall et al 2012). Typically divergence is considered a gradual process occurring via point mutations and aided by selection for original gene function plus selection for some promiscuous capacity (Bergthorsson et al 2007, Nävsall et al 2012). However, formation of chimeric proteins by gene fusion (following duplication) provides opportunity for divergence to occur rapidly and in a single - or small number - of steps. Such distal-fusion events have been reported from genomic studies (Rogers et al. 2009, Rogers and Hartl 2012) and shows how the modular nature of spatial localising domains can facilitate rapid generation of new functions (Pawson & Nash 2003).

With few exceptions (see for example studies on WspR (Bantinaki et al. 2007; Malone et al. 2007; De et al. 2008; O'Connor et al. 2012; Huangyutitham et al. 2013) and PleD (Aldridge et al. 2003; Paul et al. 2007) biochemical details underpinning the mechanisms of DGC activation and factors determining the specificity of c-di-GMP signaling are yet to be fully understood (Schirmer and Jenal 2009; Massie et al. 2012; Chou and Galperin 2016). The mechanism by which the *fadA-fwsR* fusion leads to DGC activation is unknown. The requirement for an active FwsR DGC domain and for that active domain to localize to the membrane makes clear that membrane localization is a necessary condition. One possibility is that membrane localization is alone sufficient to activate DGC activity (a suggestion bolstered by the fact that the DGC domain of *fwsR* could be fused to the membrane-spanning domain of *mwsR* with the same effect). For example, membrane localization may promote homodimerization, which is necessary for DGC activity (Wassmann et al. 2007). Activation may also be affected by fatty acid composition of the cell membrane as recently shown in *P. aeruginosa* (Blanka et al. 2015). Alternatively, localization of FwsR to the membrane may serve to facilitate spatial sequestration of c-di-GMP and thus molecular interactions between the DGC and the membrane localized cellulose biosynthetic machinery (Hengge 2009). Biochemical studies are required to explore these possibilities, but discovery that the DGC domain of FwsR becomes active upon localization opens new opportunities for understanding mechanisms of DGC activation and signaling, and points to the importance of connections between DGCs and the cell membrane. In this regard it is significant that DGCs are invariably membrane associated, or components of regulatory pathways that are - like Wsp (Bantinaki et al. 2006)) - membrane localized.

One notable feature of genes that encode DGC domains is the diverse array of domains with which they associate. Prior bioinformatic analysis of DGCs shows connections to numerous domains, ranging from CheY-like domains to kinases and phosphodiesterases (Galperin 2004; Jenal and Malone 2006; Romling et al. 2013). Gene fusion events such as we have described here are likely to contribute to the origin of these diverse spectra suggesting capacity of DGC domains to be readily accommodated within proteins of diverse function. If, as the evidence suggests, this is true, then questions arise as to why the underpinning evolutionary events have not previously been observed in experimental studies of evolution using bacteria. Our recent work provides a clue: of all single-step mutational routes to the wrinkly spreader phenotype, fusions generating chimeric proteins constitute almost 10 % of such events - but only when the genome is devoid of readily achievable loss-of-function routes to the adaptive phenotype (if loss-of-function routes are intact then fusions described here constitute ~0.1% of the total) (Lind et al. 2015). While such loss-of-function routes are readily realised in laboratory populations where functional redundancy among DGC domains is observed, the very real possibility is that in populations in natural environments such redundancy does not exist, making loss-of-function routes less achievable and mutations of the kind described here more likely. Taking experimental evolution into the wild - or at least scenarios more reflective of the complex challenges faced by natural populations (Hammerschmidt et al. 2014; Bailey and Bataillon 2016; Lind et al. 2016) is an important future goal likely to shed new light on the origins of genes.

## Materials and Methods

### Media and strains

All *P. fluorescens* strains are derivatives of *P. fluorescens* SBW25 (Silby et al. 2009) and were cultured at 28°C in King’s medium B (KB) (King et al. 1954). *Escherichia coli* DH5-a âpir (Hanahan 1983) was used to replicate all plasmids used to construct mutations. *E. coli* was cultured at 37°C in Lysogeny Broth (LB) (Bertani 1951). Solid media were prepared by the addition of 1.5% bacteriological agar. Strains intended for fluorescence microscopy were cultured in M9 minimal media (Sambrook and Russell 2001) containing 0.4% w/v glycerol and omitting thiamine. All bacterial overnight cultures were grown shaking at 160 rpm for approximately 16 hours in 30 mL glass microcosms containing 6 mL of medium. Antibiotics for the maintenance of plasmids were used at the following concentrations: gentamycin (Gm) mg L^−1^, kanamycin (Km) 100 mg L^−1^, tetracycline (Tc) 15 mg L^−1^, nitrofurantoin (NF) 100 mg L^−1^, and cycloserine 800 mg L^−1^. X-gal (5-bromo-4-chloro-3-indolyl-β-D- galactopyranoside) was used at a concentration of 60 mg L^−1^ in agar plates, IPTG at 1 mM in liquid culture and Calcofluor (fluorescent brightener 28) at a concentration of 35 mg L^−1^ in agar plates.

### Mutation reconstruction

Strand overlap extension (SOE-PCR) was employed to construct all site-directed genomic mutations in *P. fluorescens* as previously described (Ho et al. 1989; Rainey 1999; Bantinaki et al. 2007). Briefly, Phusion High-Fidelity DNA polymerase (New England Biolabs) was used to generate an amplicon product approximately 1000 bp either side of the mutation, using overlapping primer sets with the required mutation within the 5’ region of the internal primers. A second round of PCR using only the external primers was performed to make a single amplicon with the required mutation. The amplicon was then cloned into pCR8/GW/TOPO (Invitrogen) using TA cloning, transformed into an *E. coli* host and the plasmid was sequenced to confirm the presence of the mutation and no additional mutations within the amplified region. The DNA regions were then excised from pCR8/GW/TOPO by restriction digest and ligated into the pUIC3 suicide vector (Rainey 1999), and was then introduced into the *P. fluorescens* host by two-step allelic exchange (Kitten et al. 1998). This involved transconjugation of pUIC3 into the *P. fluorescens* host by tri-parental mating, and selection for transconjugants with the pUIC3 vector homologously recombined at the target sequence. Overnight non-selective cultures of the transconjugates were then treated with tetracycline in order to inhibit growth of cells that had lost the pUIC3 vector. The culture was then treated with cycloserine to kill growing cells featuring the pUIC3 vector and thus enrich the fraction of cells that were either isogenic with the host, or which now featured the mutation following a second recombination event. The culture was serially diluted and plated on KB plates containing X-gal to enable visual screening for genotypes that had lost the pUIC3 vector. Several resulting white colonies were isolated and Sanger sequencing confirmed the presence of the mutant allele and the absence of mutations that may have been introduced during the cloning process. The sequence of each genomic mutation is accessible through the following accession numbers: *fadA-fwsR* GGAAF (KU248756), *fadA-fwsR* L2M5 (KU248757), *P*_*fadA*_*-fwsR* (KU248758), *fadA-3X-fwsR* (KU248759) and *P*_*fadA*_*-mwsR1218-fwsR* (KU248760).

The visualization of proteins required translational fusions of *fadA, fwsR* and fusions of these genes with *gfp* variant *gfpmut3*\* (Andersen et al. 1998). The open reading frame of genes to be tagged were cloned into pCR8 and then moved (by digestion with restriction endonucleases and ligation) into the multiple cloning site (MCS) of mini-Tn7-LAC (Choi and Schweizer 2006). This plasmid was modified to contain both a *Pseudomonas-specific* ribosome-binding site (RBS) as well as a *gfp* encoding region 3’ of the MCS. This resulted in the ability to induce expression of our protein of interest, tagged at the C-terminus with GFP. These plasmids were sequenced to identify possible mutations introduced during cloning. Plasmids were then taken up by electrocompetent *P. fluorescens* SWB25 cells and the integration of the construct at the *attB* site was confirmed by PCR and electrophoresis. The sequence of each gfp-tagged construct cloned in the mini-Tn7-LAC MCS is accessible through the following accession numbers: *gfp* positive control (KU248761), *fwsR-gfp* (KU248762), *fadA-gfp* (KU248763), *fadA-fwsR-gfp* (KU248764) and *fadA-3X-fwsR-gfp* (KU248765).

### Microscopy techniques

Colony-level photography was performed using a Canon Powershot A640 camera in conjunction with a Zeiss Axiostar Plus light microscope using a 10x objective. Microscopy of cells was visualised using an Olympus BX61 upright microscope and images were recorded using a F-view II monochrome camera. The production of acetylated cellulosic polymer (ACP) was detected by the *in vivo* staining of colonies with Calcofluor (fluorescent brightener 28, Sigma) and this stain was visualised with fluorescence microscopy. Colonies of *P. fluorescens* were grown for 48 hours on KB plates containing Calcofluor, and several colonies were resuspended in 100 μL of distilled water. 10 μL of this solution was pipetted onto a microscope slide and used for microscopy. Stained cells were visualised using a 60x objective following DAPI excitation.

The localisation of fluorescently tagged proteins was visualised *in vivo* by fluorescent microscopy. Strains were then prepared for microscopy by inoculation of single colonies in 6 mL of sterile M9 media supplemented with 0.4% glycerol (with appropriate antibiotics to prevent loss of the mini-Tn7 vector) and grown overnight (160 rpm, 16 h). The culture was then subcultured in fresh M9-glycerol media (without antibiotics) resulting in an OD_600_ ~0.05 culture. This was then incubated with shaking for 60 minutes, after which 1 mM Isopropyl p-D-1-thiogalactopyranoside (IPTG) was added to induce expression of the tagged genes. The culture was then returned to the incubator for 2 hours until an OD_600_ of approximately 0.3 was reached. In order to increase the density of cells for microscopy, a 1 mL aliquot of culture was centrifuged and resuspended in 100 μL of M9-glycerol media. Agarose pads were prepared consisting of M9-glycerol media with 1% w/v agarose. A 3 μL aliquot of resuspended culture of induced cells was pipetted onto the pads, which were left to dry and then covered with a coverslip. Cells were visualised using an Olympus BX61 upright microscope and both phase contrast and fluorescence images were recorded using a F-view II monochrome camera. Fluorescence images were observed using a constant 3500 ms exposure time across all strains. All images were processed using FIJI (Schindelin et al. 2012) to obtain measures of fluorescence and measures of co-localisation. Measures of fluorescence levels were obtained by calculating the mean gray values of cell transects observed under fluorescence microscopy. Four randomly picked cells were analyzed across two independent images for each biological replicate, with three biological replicates analyzed per genotype. Co-localisation analyses were prepared by generating the mean van Steensel’s distribution of two fields of view for each biological replicate. At least three biological replicates were used over two separate experiments in co-localisation analyses.

## Acknowledgements

This work was supported in part by Marsden Fund Council from New Zealand Government funding (administered by the Royal Society of New Zealand) and the New Zealand Institute for Advanced Study. We thank Jenna Gallie and Peter Lind for discussion and comments on the manuscript and Heather Hendrickson for assistance with microscopy.

## Figure Supplements

**Figure 1 - figure supplement 1:**
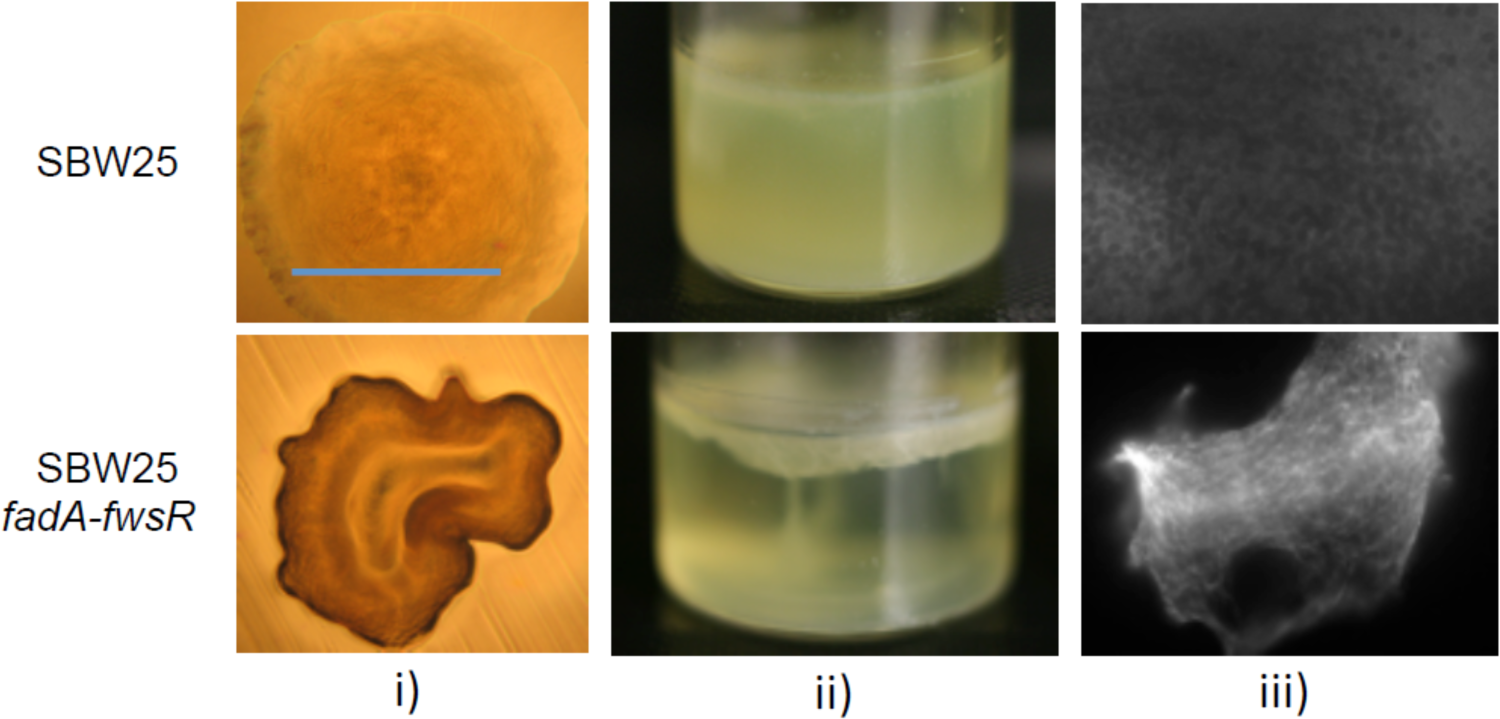
The reconstruction of the *fadA-fwsR* mutation in ancestral backgrounds causes WS. Genotypes featuring *fadA-fwsR* (either naturally evolved or reconstructed in SM backgrounds) are characteristically WS in terms of i) morphology (visualised by light microscopy (10x objective) of ~32 h colonies grown on King’s Medium B agar (KB), blue bar is ~2mm), ii) formation of mats at the air-liquid interface of microcosms (incubated statically at 28^o^C for 3 days) and iii) calcofluor binding as indicative of ACP biosynthesis (visualised by fluorescent microscopy, 60x objective). SBW25 is ancestral *P. fluorescens* SBW25, SBW25 *fadA-fwsR* has the causal *fadA-fwsR* mutation reconstructed into the ancestral SM background.

**Figure 1 - figure supplement 2:**
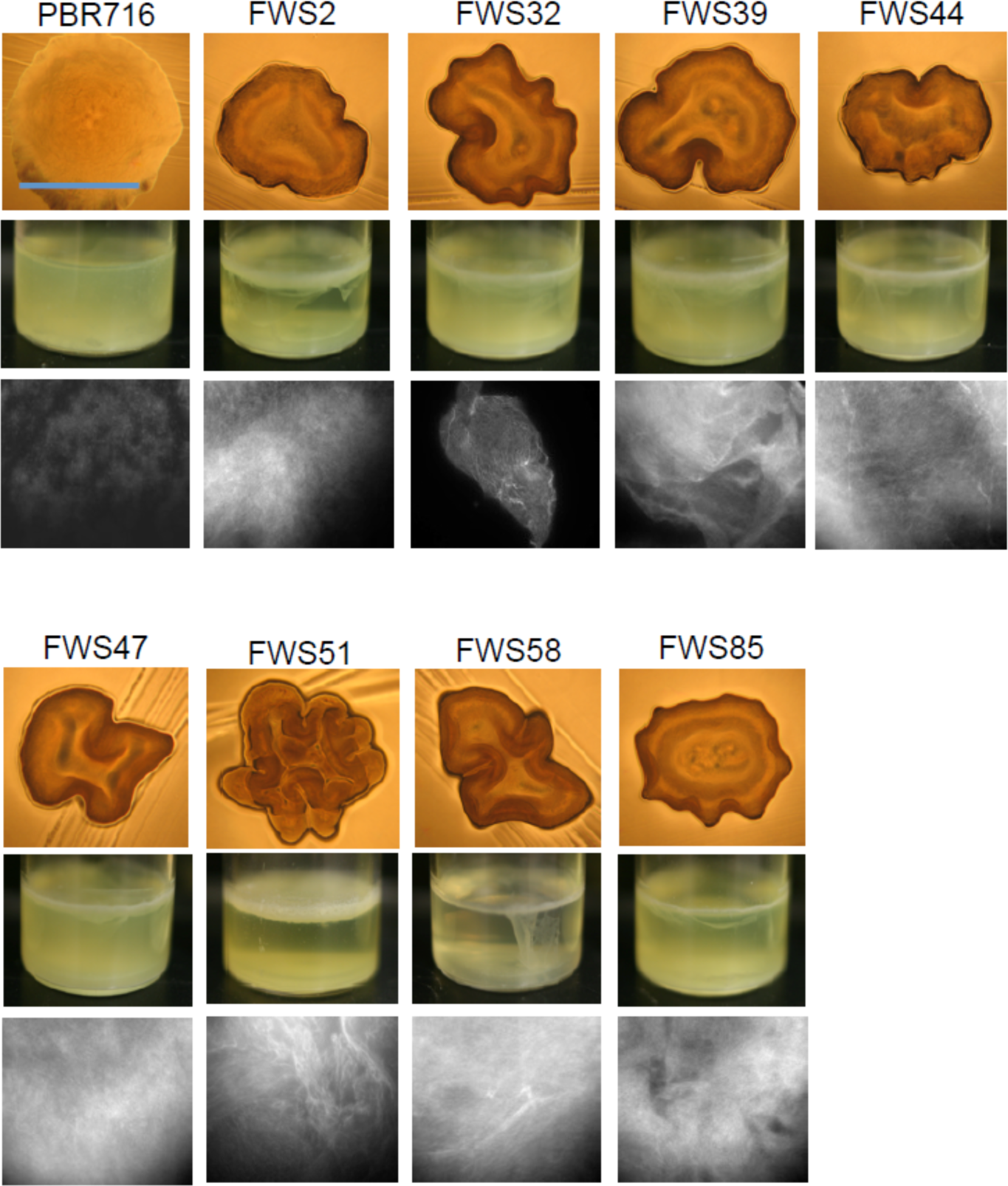
The spectrum of *fadA-fwsR* fusions arising from SBW25 Δ *wsp aws mws* (PBR716) express the WS phenotype. Numbered are the different isolates with *fadA-fwsR* fusion mutations. The phenotypes of the *fadA-fwsR* fusions are characteristic WS in morphology, colonisation of the air-liquid interface and ACP production (see Figure 1 - figure supplement 1 for details of each image).

**Figure 1 - figure supplement 3:**
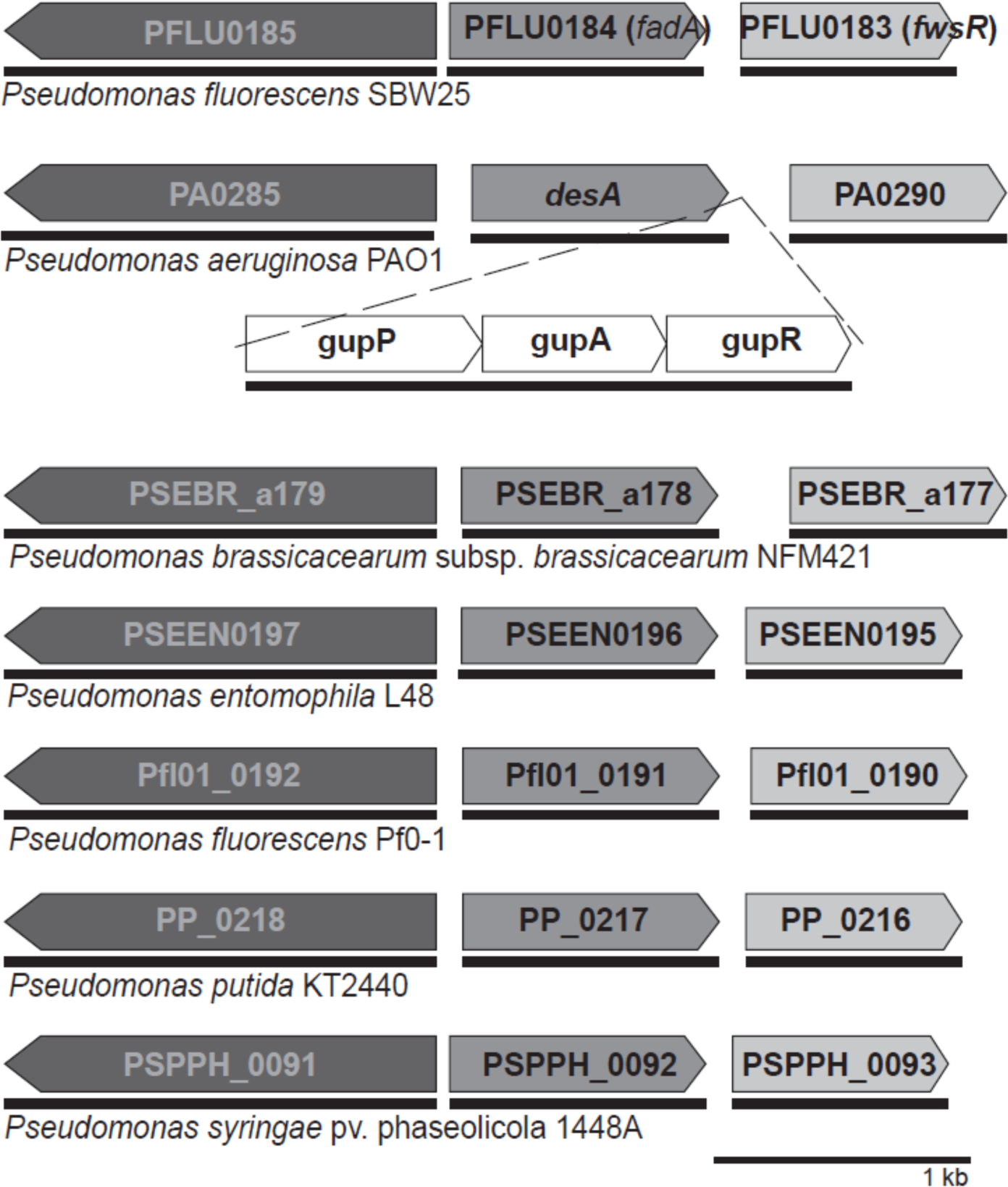
Organisation of homologs of *fadA* and *fwsR* across *Pseudomonas* spp. The arrangement of neighbouring genes of the *fwsR* locus is conserved across many species of the genus *Pseudomonas.* Notable exceptions are strains of *P. aeruginosa*, in which the paralogs *fadA* and *fwsR* are separated by a single predicted operon of genes *gupPAR.* There is no evidence of recombination between *fadA* and *fwsR* in other strains. Genes sharing the same shade represent likely paralogs as determined by the *Ortholuge* web program (Whiteside et al. 2013), with thin black bars (underneath depicted ORFs) representing operons as predicted by the *DOOR* web program (Mao et al. 2014)

**Figure 7 - figure supplement 1:**
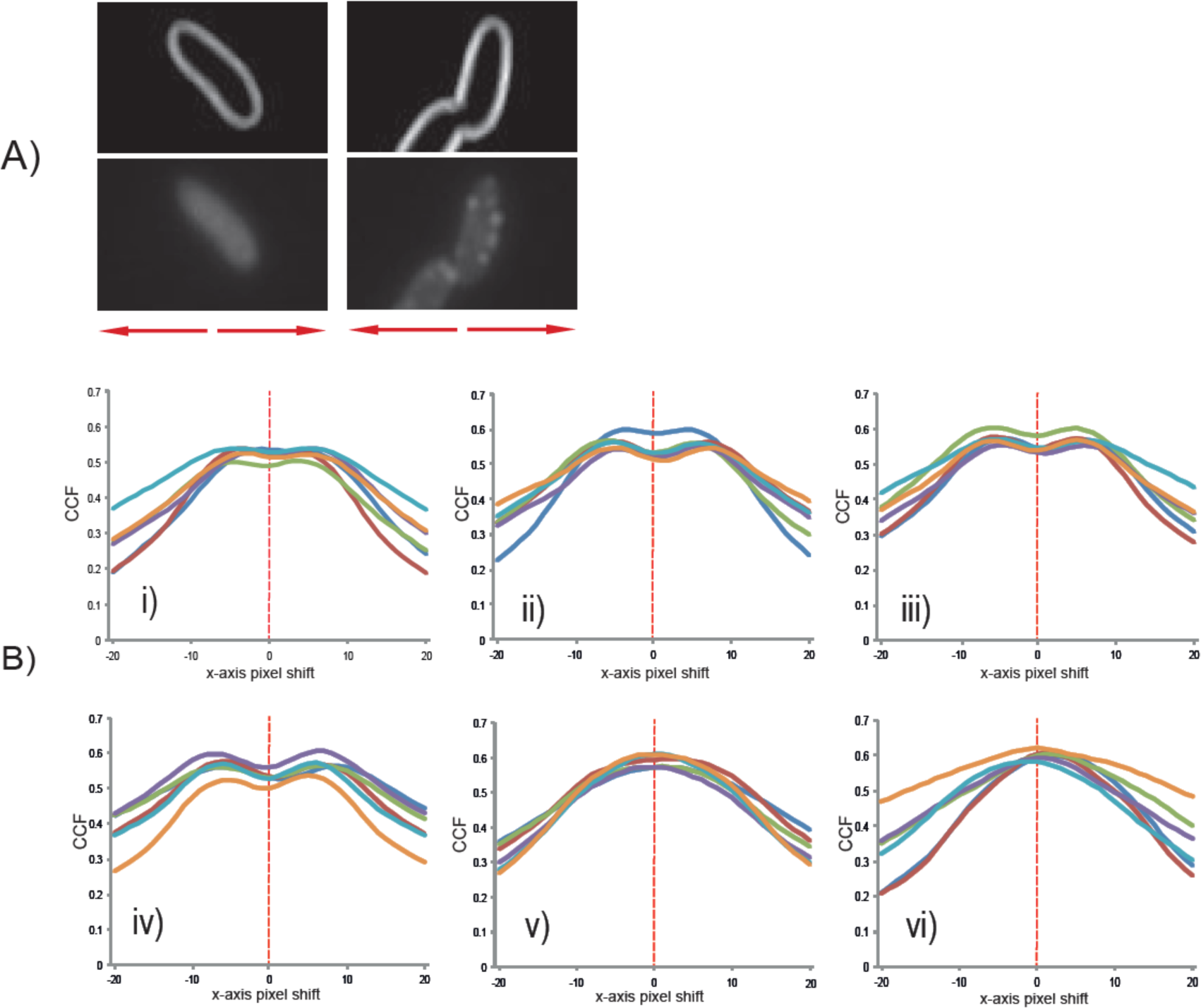
van Steensel’s distribution of colocalisation of fluorescent signal and the cell edge. **A)** The CCF represents the degree of colocalisation of the reference image (the marked edges of cells observed by phase contrast microscopy) and the fluorescent distribution of images. The edges of cells were marked using FIJI (Schindelin et al. 2012). Depicted are representative images of cells expressing *fwsR-gfp* (left) and *fadA-fwsR-gfp* (right). To generate van Steensel’s distributions, the images are moved horizontally relative to each other and the CCF value is calculated using FIJI with the JACoP plugin (Bolte and Cordelieres 2006). **B)** Graphs represent van Steensel’s distributions for i) gfp(-) control; ii) gfp(+) control; iii) *fwsR-gfp;* iv) *fadA-3X-fwsR-gfp;* v) *fadA-gfp;* and vi) *fadA-fwsR-gfp.* Graphs i, ii, iii and iv depict bimodal distributions indicative of a fluorescent distribution in between the reference markings of the cell edge. Distributions v and vi are unimodal and the distribution peaks approximately at a pixel-shift of 0, indicative of direct correlation of the fluorescent signal with the cell edge. Analyses of six biological replicates were performed per genotype, with each replicate the analysis of approximately 100 cells analysed per biological replicate.

